# Multi-omics analysis of mouse fecal microbiome reveals supplier-dependent functional differences and novel metagenome-assembled genomes

**DOI:** 10.1101/2022.09.12.507288

**Authors:** Zachary L McAdams, Susheel Bhanu Busi, Kevin L Gustafson, Nathan Bivens, Craig L Franklin, Paul Wilmes, Aaron C Ericsson

**Author notes:** These authors contributed equally to this project (Zachary L McAdams, Susheel Bhanu Busi). **Corresponding author:** ACE.

## Abstract

Host genetics, sex, and other within-source factors have been associated with characteristic effects on the fecal microbiome in mice, however, the commercial source of mice remains the dominant factor. Increasing evidence indicates that supplier-specific microbiomes in particular confer differences in disease susceptibility in models of inflammatory conditions, as well as baseline behavior and body morphology. However, current knowledge regarding the compositional differences between suppliers is based on 16S rRNA amplicon sequencing data, and functional differences between these communities remain poorly defined. Here, we applied a meta-omic (metagenomic and metatranscriptomic) approach to biomolecules (DNA/RNA) extracted from murine fecal samples representative of two large U.S. suppliers of research mice, which differ in composition, and influence baseline physiology and behavior as well as disease severity in mouse models of intestinal disease. We reconstructed high-quality metagenome-assembled genomes (MAGs), frequently containing genomic content unique to each supplier. These differences were observed both within pangenomes of dominant taxa as well as the epibiont *Saccharimonadaceae*. Additionally, transcriptional activity and pathway analyses revealed key functional differences between the metagenomes associated with each supplier, including differences in carbohydrate enzyme activity and dissimilatory sulfate reduction by sulfate-reducing bacteria (SRB). These data provide a detailed characterization of the baseline differences in the fecal metagenome of laboratory mice from two U.S. commercial suppliers suggesting that these functional differences are influenced by differences in the initial inoculum of colony founders, as well as additional taxa gained during growth of the production colony.

## Introduction

Host-associated microbiomes, such as the gut microbiome (GM), exert strong effects on host physiology, susceptibility or resistance to various conditions, and response to treatment and dietary challenges. Investigations at the population-level suggest that differences in the human GM are responsible for a large portion of the variability within individual host responses to a given dietary challenge^1,2^ or medical treatment,^3–5^ implying that the GM is an important consideration in precision health and medicine strategies. Similarly, the GM of laboratory mice within the biomedical research community is highly variable due to numerous covariates,^6,7^ and these compositional differences have been associated with differences in host fitness in the context of uniform host genetics and environment.^8–12^ One of the dominant factors contributing to the population-level variability in specific pathogen-free (SPF) mouse microbiomes is the commercial source of mice.^13–15^ Previous studies have demonstrated reproducible differences in the GM richness and beta-diversity, irrespective of host genotype (i.e., strain) within each supplier.^12,13^ Specifically, the GM of mice supplied by the Jackson Laboratory (Jackson) and Envigo are characteristically of low and high richness, relative to each other, and each comprises unique taxa, in addition to an apparent core population of bacteria common to both sources. The latter includes members of the semi-standardized altered Schaedler flora (ASF),^16,17^ reflecting the common procedures used to establish mouse production colonies on a commercial scale. Suppliers often surgically transfer embryos of the desired genotype to a pseudopregnant surrogate dam colonized with ASF, which then seeds the initial generation of offspring with that limited microbiome comprising 8 to 10 cultivable bacteria.^18^ These mice are then used to establish multiple generations of filial mating to expand the colony, during which time mice are housed in large open-top caging systems and allowed to become colonized with additional bacteria from the environment. It is believed that subtle environmental differences are responsible for the reported supplier-origin differences, as well as the differences between multiple distribution facilities of the same supplier^13^ or changes within a supplier over time.^19,20^

However, GMs with different taxonomic compositions may possess qualitatively similar functional capacities.^21,22^ It is therefore unclear whether the different GMs colonizing mice from Jackson and Envigo are functionally different. Owing to the reported influence of these GMs in multiple mouse models of disease,^23–25^ we hypothesized that the compositional differences result in substantial functional differences, as evaluated by the metatranscriptome. Any detected differences in the functional capacity of the fecal microbiome could therefore be due to differences in the ASF isolates maintained by each institution, the environmental exposures during colony expansion, or both. As researchers continue to leverage the inherent differential effects of these complex GMs as a population-level model of human host/microbe interactions, it is important to understand the differences in the metagenome and transcriptional activity of mice from these different suppliers, and the origin of any detected differences. With those goals in mind, fecal samples from healthy adult CD-1 mice colonized with a Jackson-origin or Envigo-origin GM (GM1 and GM4, respectively) were collected and used as the source of DNA and RNA for metagenomic and metatranscriptomic analyses using a reiterative co-assembly procedure. We report here the identification of 86 high- and medium-quality novel and previously identified metagenome-assembled genomes (MAGs), analyzed and compared in the context of the two source GMs, and a detailed description of the functional differences between mice from these two commercial sources of SPF mice.

## Results

### Metagenomic, metatranscriptomic and taxonomic summary

To establish a taxonomic and functional profile, using IMP^26^ (v3, commit #9672c874; available at https://git-r3lab.uni.lu/IMP/imp3) 2.09 × 10^9^ metagenomic and metatranscriptomic reads were co-assembled and binned into MAGs. Subsequently, the completeness and contamination were assessed using CheckM. Per established criteria in the field,^27^ **Table 1** lists the 29 high-quality (>90% completion and < 5% contamination) MAGs from the entire dataset (**Figure 1A**). An additional 35 medium-quality (> 80% completion and <10% contamination) MAGs, 22 medium-quality MAGs with completeness >50%, 17 low-quality (partial) MAGs with between 31% and 49% completeness, and 25 MAGs with >50% completeness but >10% contamination were identified (**Figure 1A)**. A complete list of the 128 identified MAGs is provided as **Supplementary Table 1**. Over 75% of MAGs contained greater than 20 tRNA-encoding genes, with over half encoding 30 or more tRNA genes (**Figure 1B**). Over 75% of the 128 assembled MAGs had an average coverage of 10× or greater (median 24.4×, range 2.1× to 1540×; **Supplementary Table 1**) and roughly 65% of MAGs (including the majority of high-quality MAGS) were assembled from less than 200 scaffolds (**Figure 1C**). Comparison of metagenomic composition and the metatranscriptome revealed a strong correlation, suggesting transcriptional activity of the majority of detected genes (**Figure 1D**). As expected, there was also a strong correlation between the number of genes detected and total size of the assembled MAGs (**Supplementary Figure 1**).

**Figure 1.**
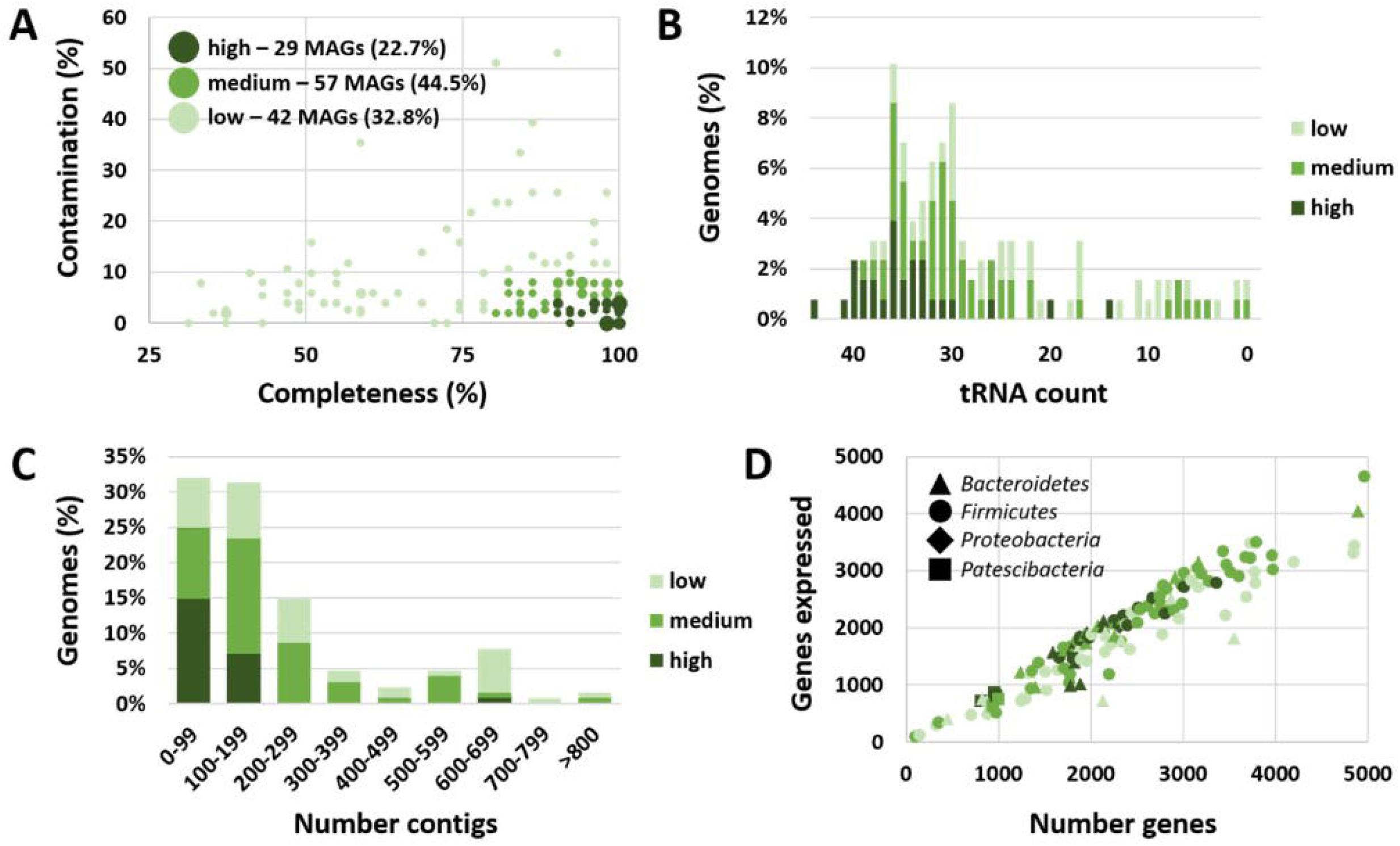
Dot plot showing the completeness (%) and contamination (%) among the 128 metagenome-assembled genomes (MAGs) recovered from all six samples, legend in inset, dot size correlates to the number (1 to 6) MAGs represented (**A**); bar charts showing the number of tRNAs found in low-, medium-, and high-quality MAGs (**B**), and the number of contigs used to construct MAGs (**C**); and dot plot showing the number of expressed genes in relation to total detected genes in each MAG (**D**).

Of the 64 high- and medium-quality MAGs with > 80% completion and <10% contamination listed in **Supplementary Table 1**, over one third (23/64) were identified as members of the Gram-positive family *Lachnospiraceae* (phylum *Bacillota*). The second most common family was the Gram-negative *Muribaculaceae*, within the phylum *Bacteroidota*. Other MAGs within the phylum *Bacillota* included several members of the *Ruminococcaceae, Clostridiacae, Bacillaceae*, and *Lactobacillaceae*, among others. Additional MAGs within the *Bacteroidota* included members of the genera *Alistipes, Bacteroides, Parabacteroides, Odoribacter*, and *Prevotella*. Six of the high- and medium-quality 64 MAGs in **Supplementary Table 1** were external to either of those two dominant phyla, including one identified as *Parasutterella excrementihominis* (*Burkholderiaceae* within phylum *Pseudomonadota*), and five identified as members of the family *Saccharimonadaceae* (phylum *Patescibacteria*).

### Candidate Phyla Radiation taxa demonstrate strain-level differences between vendors

As newly recognized epibionts within the candidate phylum radiation (CPR), the *Saccharimonadaceae* were of particular interest since their reports in laboratory mouse strains are limited. MAGs identified as *Saccharimonadaceae* have been found in diverse environmental samples including deep sea hydrothermal vents, glacial-fed stream biofilms,^28^ and petrochemical plant sludge.^29^ Regarding host-associated samples, *Saccharimonadaceae* are most commonly identified in human oral cavity samples,^30,31^ although a handful of rumen^32^ and fecal^33^ samples have also yielded MAGs. A thorough search of the National Center for Biotechnology Information (NCBI) Sequence Read Archive (SRA) found 321 MAGs within this phylum, including four MAGs from mouse feces. Comparison of the phylogenetic relationship of the newly generated five MAGs within *Saccharimonadaceae* revealed similarity to other host-associated isolates, and particularly the mouse-origin MAGs (**Figure 2A**). Construction of a *Saccharimonadaceae* pangenome from the current data revealed portions of highly conserved core genomic content, and regions of genomic material specific to MAGs from either of the two supplier-dependent microbiomes (**Figure 2B**), suggesting the vendors each harbor distinct strains of this taxonomy, with distinct functional capacities. These data also suggested the co-evolution and transfer of genetic material between bacterial strains within each source.

**Figure 2.**
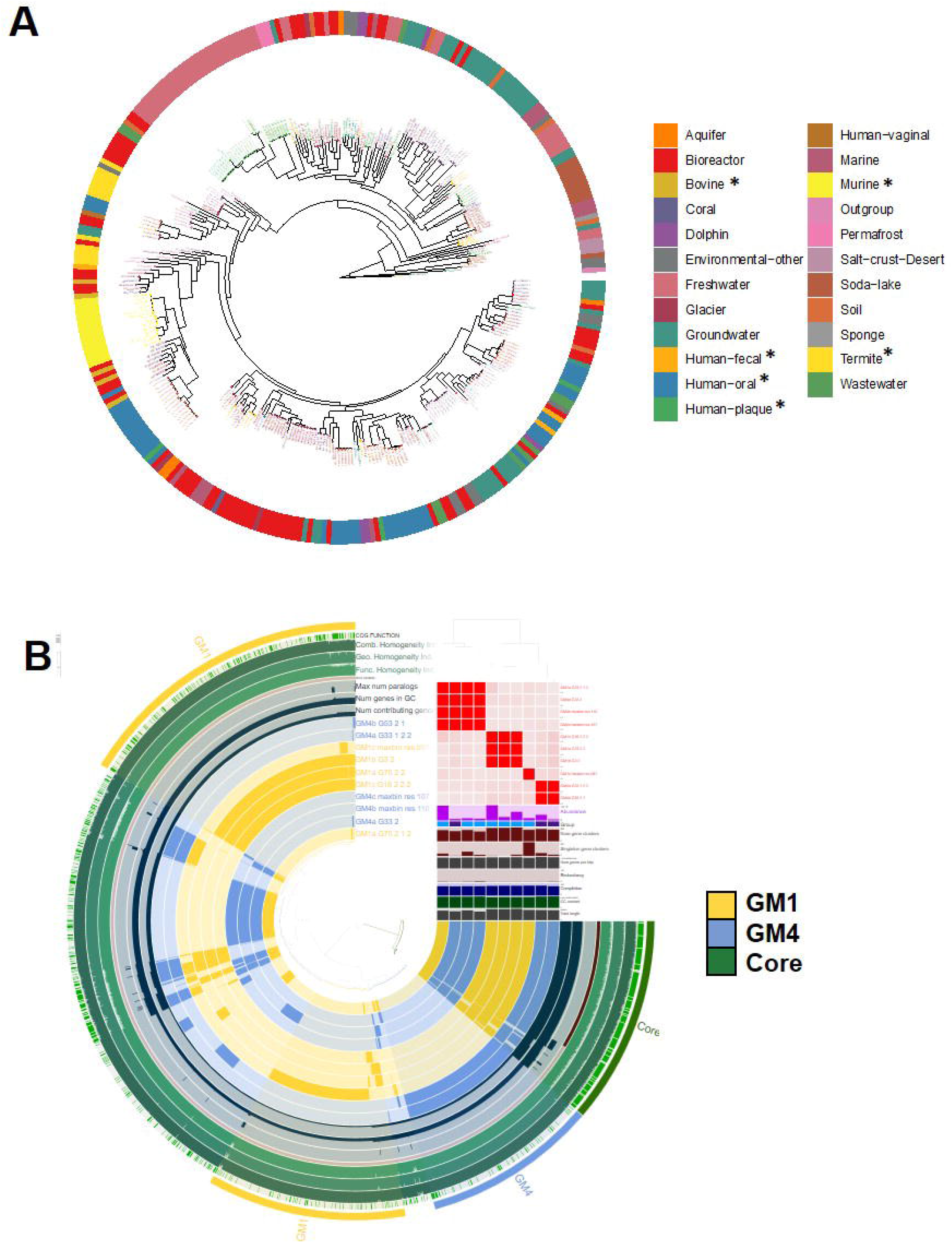
Phylogenetic tree (ignoring branch lengths) showing the relationship between the newly identified *Saccharimonadaceae* MAGs and 321 MAGs within the NCBI Sequence Read Archive (SRA) annotated to the *Saccharimonadaceae* family, asterisk represents gut-associated samples (**A**); and pangenome of novel *Saccharimonadaceae* MAGs showing genomic content specific to MAGs from each source (**B**).

### Distinct source-dependent MAGs within multiple taxonomies

To further investigate the genomic heterogeneity within other common taxonomies, separate pangenomes were created for various members of the Gram-negative phylum *Bacteroidota*, including *Alistipes* spp. (10 MAGs, **Figure 3A**), *Prevotella* spp. (9 MAGs, **Figure 3B**), and family *Muribaculaceae* (17 MAGs, **Figure 3C**). As in the *Saccharimonadaceae* pangenome comparison, each genus or family revealed regions of genomic content conserved between multiple MAGs from each of the supplier-origin microbiomes, along with conserved core genomic content encoding for single copy gene (SCG) clusters, suggesting that the transfer of genetic material is an ongoing process within each of these taxonomies, at each production source. Similarly, pangenomes were constructed from dominant members of the Gram-positive phylum *Bacillota*, including *Lactobacillus* spp. (14 MAGs, **Figure 4A**) and family *Lachnospiraceae* (20 MAGs, **Figure 4B**). These pangenomes revealed conserved genomic content including highly conserved SCG clusters within each taxonomy, as well as source-dependent differences in the genomic content of MAGs, which may be interpreted as evidence of distinct lineages of Gram-positive taxonomies in mice from each supplier. Collectively, these data indicate the presence of substantial differences in the functional capacity of the dominant bacterial families detected in the microbiome of mice from different suppliers.

**Figure 3.**
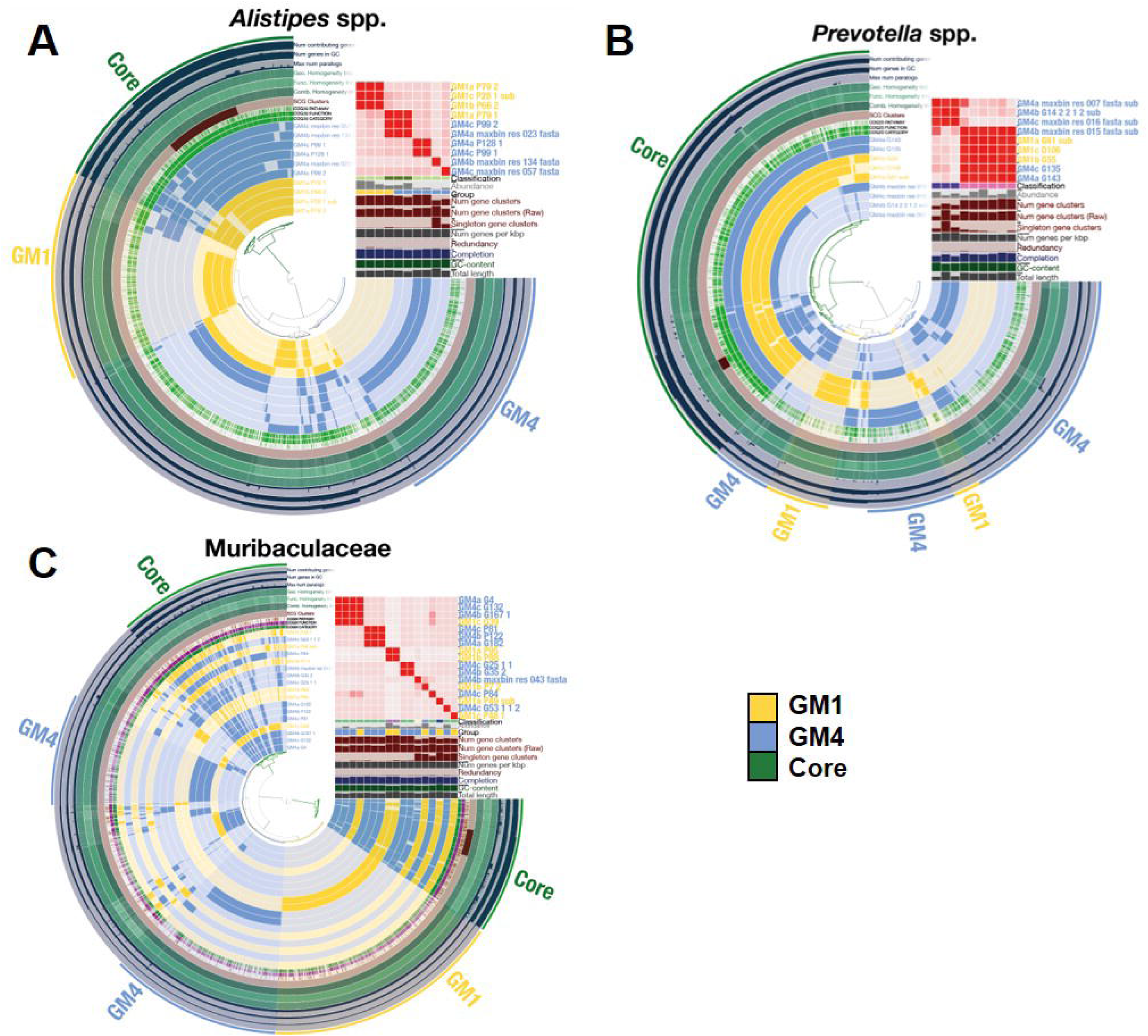
Pangenomes of *Alistipes* (**A**), *Prevotella* (**B**), and family *Muribaculaceae* (**C**) constructed from the present data, each showing the conserved core genomic content, and additional genomic content, common to multiple MAGs from each supplier

**Figure 4.**
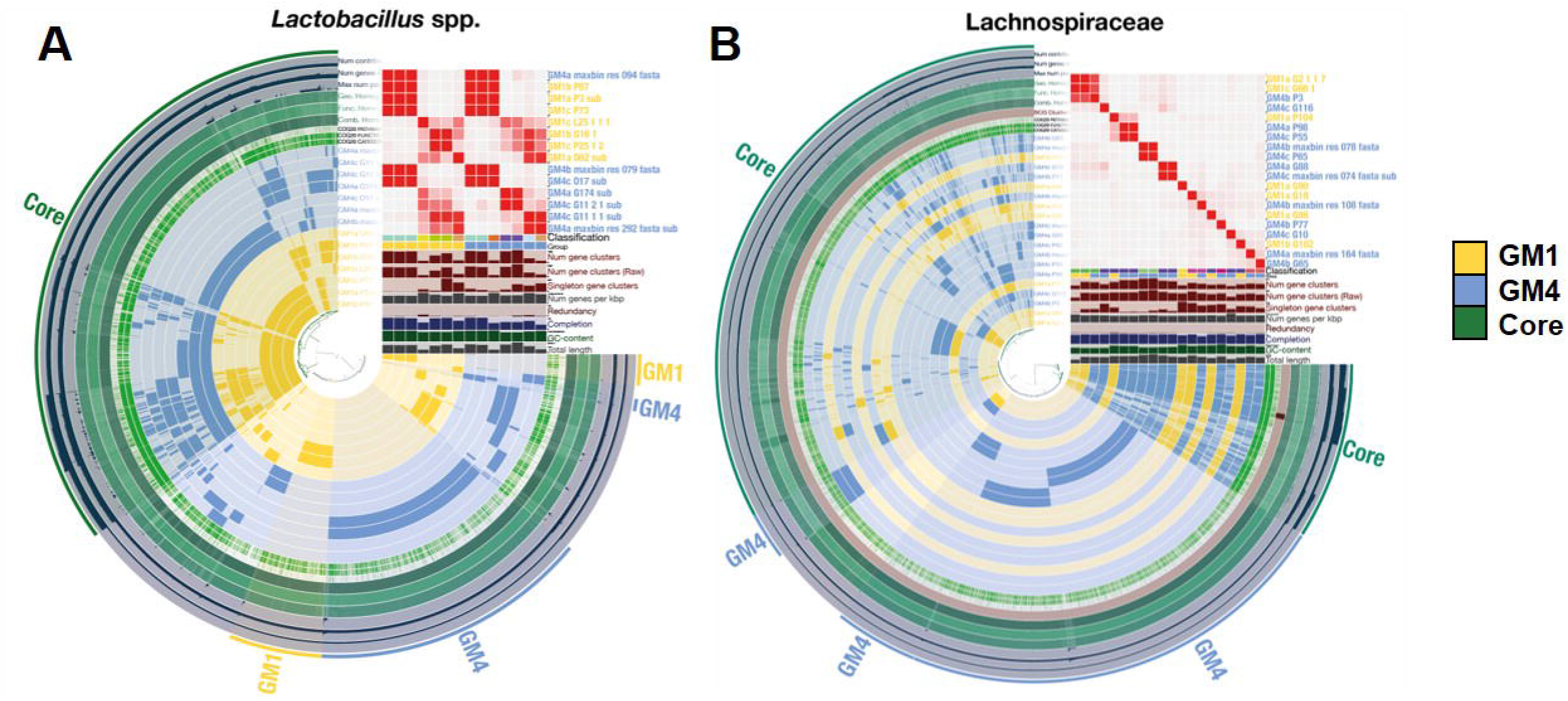
Pangenome of *Lactobacillus* (**A**) and family *Lachnospiraceae* (**B**) constructed from the present data, each showing the conserved core genomic content, and additional genomic content, common to multiple MAGs from each supplier.

### Functional differences between source-dependent GM

Based on the above observations and our original hypothesis, the metatranscriptome was compared between GM profiles to determine if the detected differences in metagenomic content were also associated with differences in transcriptional activity. Transcripts were compared to (and cross-referenced against) multiple databases including the Kyoto Encyclopedia of Genes and Genomes (KEGG),^34,35^ the Protein family (Pfam) database,^36^ and the CAZy database^37^ of carbohydrate active enzymes and accessory molecules. **Figure 5A** shows KEGG-identified microbial-associated pathways comprising a multitude of differentially expressed KEGG orthologs (**Figure 5B**). A list of differentially expressed KEGG-identified host-associated pathways is shown in **Supplementary Figure 2**. Similarly, comparison of the bulk metatranscriptome annotations against the Pfam (**Figure 5C**) and CAZy (**Figure 5D**) databases resulted in many differences, with greater transcriptional activity of different genes in each GM. **Supplementary File 1** lists all significantly differing KEGG, Pfam, and CAZyme annotations as determined by DESeq2^38^ (*p* < 0.05).

**Figure 5.**
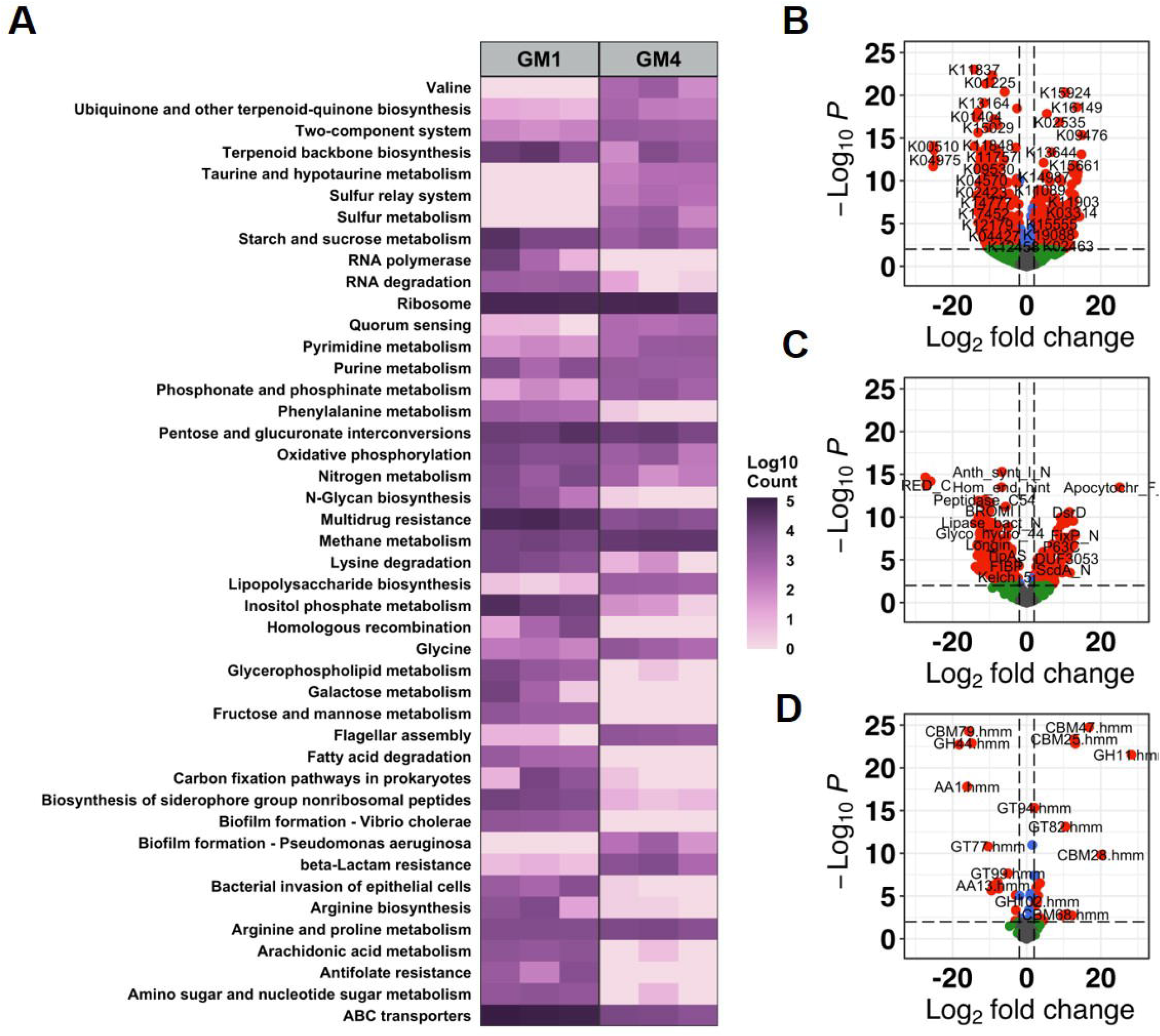
Heat map of differentially expressed select KEGG pathways (A) and volcano plots of individual KEGG (B), Pfam (C), and CAZyme (D) IDs between Jackson (GM1)- and Envigo (GM4)-origin microbiomes. Differentially abundance testing was performed using DESeq2 with *p* < 0.05 considered significant.

Source-dependent differences within the KEGG annotations included several GM1- and GM4-specific genes involved in a wide range of functions. To increase our ability to discern biologically meaningful differences in the function of these GMs, the top 25% most significant KEGG IDs (lowest p values identified by DESeq2) found to significantly differ between GM1 and GM4 were manually reviewed and curated to identify multiple KEGG IDs within a pathway, and thus likely representing true differences in the functional activity of that pathway (**Figure 5A-B**). Several GM1-specific genes involved in diverse metabolic functions were identified including starch and sucrose (CBH1, K01225; SI, K01203), and fructose and mannose (algG, K01795; mtlK, K00045), arachidonic acid (EPHX2, K08726), and phenylalanine (mhpF, K04073; DDC, K1593) metabolism.

Source-dependent differences within the KEGG annotations also included several GM4-specific genes involved in numerous functions including flagellar assembly (flgH, K02393; flgI, K02394, flgA, K02386), quorum sensing (srfATE, K15657), lipopolysaccharide biosynthesis (lpxC, K02535; lpxI, K09949), and sulfur metabolism (dsrA, K11180). Pfam annotations also identified increased expression of genes within the dissimilatory sulfite reduction pathway (DsrC, DsrD, and FdhE) and chemotactic responses (CheZ, TarH) by bacteria within GM4. Among the many genes found to be differentially expressed, patterns emerged suggesting increased activity of certain pathways in GM4, including the tricarboxylic acid (TCA) cycle and cytochrome c oxidase activity. Increased TCA cycle activity is suggested by increased expression of enzymes within the TCA cycle (succinate dehydrogenase D; sdhD); enzymes involved in acetyl-CoA production (malonyl-CoA/succinyl-CoA reductase; mcr); three different ccb-type cytochrome c oxidase subunits (I, II, and III) and the fixS cytochrome c oxidase maturation protein; and the cytochrome c-type biogenic protein ccmE. Additionally, GM4 had increased expression of enzymes associated with acetate (acetoacetate decarboxylase, adc), propanoate (methylmalonyl-CoA mutase, mcmA1), and butanoate production (mcr) using TCA cycle compounds, suggesting that the increased release of stored energy by the GM may be associated with increased production of compounds beneficial to the host such as short chain fatty acids (SCFAs).

Lastly, numerous source-dependent differences in carbohydrate active enzymes (CAZymes) and accessory molecules were identified (**Figure 5D)**. The glycoside hydrolase (GH) family 48 (GH48.hmm) including chitinase and cellulobiohydrolases enzymes was differentially expressed in GM1 using both CAZyme and Pfam (Glyco_hydro_48) annotations. Other GM1-associated CAZyme molecules included the auxiliary activities (AA) of multicopper oxidases (AA1.hmm) and glycosyltransferase (GT) families that bind the LPS inner core polysaccharide^39^ (GT99.hmm) and the host-produced extracellular polysaccharide heparan^40^ (GT64.hmm). GM4-associated CAZymes included multiple non-catalytic carbohydrate binding motifs (CBMs) with diverse targets including β-1,3-glucan and LPS (CBM39.hmm), cyclodextrins (CBM20.hmm), lactose (CBM71.hmm), and fucose (CBM47.hmm). CBMs specific to cellulose and chitin were identified in both GM1 (CBM2.hmm, CBM72.hmm) and GM4 (CBM28.hmm). Collectively, these data demonstrate extensive differences in the baseline transcriptional activity at the enzyme and pathway levels of supplier-origin gut microbiomes.

### Supplier-origin GMs indicate variable levels of enzymatic activity associated with eukaryotes

While most studies focus on bacterial abundances and differences, we observed eukaryotic organisms present within each GM (*Methods)*. The largest portion of eukaryotes identified belonged to the phylum *Ochrophyta* followed by *Dinoflagellata* and *Chlorophyta*, all within the kingdom Protista **(Supplemental Figure 3)**. Eukaryotes identified within the kingdom Fungi were constrained to the phylum *Ascomycota* with limited taxonomic resolution. No significant differences in the relative abundance of eukaryotic phyla were observed between GMs. Interestingly, while no differences in the relative abundance of eukaryotic phyla were observed between GM1 and GM4 (**Supplemental Figure 3**), glycoside hydrolase CAZyme expression was negatively correlated with GM1 eukaryotes while positively correlated to GM4 eukaryotes (**Supplemental Figure 4**). Lastly, we identified several genes with increased expression in GM1 previously associated with a wide range of host metabolism and disease pathways (**Supplementary Figure 2**), however, the biological significance of the differential expression of these pathways remains unclear.

## Discussion

The Jackson (GM1)- and Envigo (GM4)-origin GMs influence many host phenotypes including intestinal inflammation,^23^ colonization resistance,^25^ and behavior and body morphology.^24^ The robust taxonomic differences between these supplier-origin GMs influencing phenotypic differences have previously been identified using targeted amplicon (e.g., 16S rRNA) sequencing, however, this approach yields limited taxonomic resolution of detected amplicons, and a complete lack of information regarding functional capacity or transcriptional activity. Using an iterative co-assembly procedure, we combined metagenomic and metatranscriptomic sequencing of the fecal microbiome of laboratory mice to provide a valuable resource describing the baseline metagenomic and transcriptional differences of Jackson- and Envigo-origin GMs (GM1 and GM4, respectively). The current data build upon earlier reports of differences in the composition of the GM in mice from different suppliers^25,41,42^ by providing a more detailed assessment of those differences as well as functional differences.

Many of those functional differences were attributable to differences in bacteria associated with the ancestral ASF used in the colony founders, including *Lactobacillus murinus* [ASF361] and *L. intestinalis* [ASF360]. These differences could therefore ostensibly be controlled or changed during the initiation of new production colonies. An additional notable aspect of the source-dependent differences in *Lactobacillus* function is the growing body of evidence supporting *Lactobacillus* spp. as psychobiotics,^43^ or live organisms capable of conferring benefits to mental health when ingested. Recent studies have demonstrated differences in anxiety-related behavior and spontaneous locomotor and exploratory behavior between isogenic mice harboring GM1 or GM4^24^ and *L. intestinalis* and related species have both been shown to confer vagus nerve-dependent effects on behavior.^44–46^ Differences in the genomic content of these MAGs provides one possible explanation for the host phenotypic differences.

Numerous differentially expressed KEGG orthologs representing several microbial- and host-associated pathways were identified between the supplier-origin GMs **(Figure 5A-B, Supplemental Figure 2)**. Consistent with the previously reported differences in *Pseudomonadota*^13^ of GM1 relative to GM4, the Jackson-origin GM was associated with decreased expression of lipopolysaccharide biosynthesis and flagellar assembly. Low richness microbiomes have been associated with increased body weight, growth,^47^ and intestinal inflammation.^23^ Here we have identified that, in addition to fatty acid degradation, multiple carbohydrate metabolic pathways including starch and sucrose, galactose, and fructose metabolism were increased in the low-richness GM. The differential expression of these metabolic pathways may increase energy availability to the host likely contributing to the GM1-associated increase in body weight and growth^24^ and increased intestinal inflammation in models of intestinal disease.^10,48,49^

A differentially expressed KEGG pathway in GM4 that can be linked to previously recognized compositional differences is the dissimilatory sulfite reduction (DSR) pathway, expressed by sulfate-reducing bacteria (SRB) such as *Desulfovibrio* and *Bilophila* spp. These taxa, identified as unique GM4-associated features^50^ are responsible for production of H_2_S, a compound with biphasic effects on inflammation, hypertension, and tumorigenesis depending on its intra- and extracellular concentrations.^51–57^ Thus, augmentation of intracellular H_2_S production by luminal SRB may result in the low levels adequate to confer protective effects in certain scenarios, or sufficiently high to adversely influence disease susceptibility in others.

These data are also of interest from an evolutionary perspective, as they provide a glimpse of the short-term evolutionary landscape within the GM at each supplier. Pathogenic bacteria frequently undergo rapid evolution within a host organism through recombination and mutation,^58,59^ and the same events occur between and within commensal members of the microbiota.^60,61^ Moreover, pathobionts can arise from commensal organisms through the same mechanisms.^62^ In the data presented here, the consistent finding of source-specific genomic content within genera suggests separate evolutionary trajectories at each supplier, occurring in all dominant taxonomies with multiple closely related members. Notably, this feature was particularly evident in the relatively small pangenome of *Saccharimonadaceae*. These findings are in agreement with the recent study from Yilmaz et al. demonstrating the long-term evolution of microbiota and development of multiple co-existing substrains of bacteria within individual taxonomies.^63^

Lastly, we were surprised to recover a large number of high-quality MAGs associated with the family *Saccharimonadaceae* (formerly known as TM7), epibionts^64,65^ which were unrecognized until their identification using molecular methods. Successful culture requires co-culture with the cognate host bacteria, including *Actinomyces odontolyticus* and other members of the human oral cavity.^65,66^ That being said, these highly auxotrophic epibionts with extremely limited genomes are found in virtually all environmental conditions while being surprisingly scarce in metagenomic data from fecal microbiomes. Our analysis agrees with that by Dinis *et al*.,^67^ which demonstrated that the vast majority of host-associated MAGs from this phylum were from human oral cavity samples or rumen contents, with much fewer fecal samples represented. It is unclear which bacteria serve as the host for fecal members of *Saccharibacteria*.

Detecting differences in microbial diversity and composition between Jackson- and Envigo-origin GMs has previously relied upon targeted amplicon sequencing of the 16S rRNA gene.^13,24^ While informative, this approach is limited by taxonomic resolution and does not provide the baseline functional capacity or transcriptional activity of these GMs. Our metagenomic and metatranscriptomic sequencing of Jackson- and Envigo origin GMs has established that distinct differences in both the functional capacity and baseline transcriptional activity at the gene and metabolic pathway levels exist amongst the dominant taxa within supplier-origin GMs. Collectively, these data will serve as a valuable resource to leverage the host-microbiome relationship in mouse models of disease and behavior in future.

## Methods

### Mice and sample collection

Mice contributing fecal samples were eight-week-old, female, CD-1 mice produced by breeding colonies maintained at the MU Mutant Mouse Resource and Research Center in accordance with the Guide for the Care and Use of Laboratory Animals approved by the University of Missouri Institutional Animal Care and Use Committee (IACUC, protocol 9587). Mice colonized with a Jackson-origin GM (GM1) or Envigo-origin GM (GM4)^42^ were housed in microisolator polycarbonate cages on individually ventilated racks, under positive pressure. A sample size of three mice per GM was selected to attain a power of 80% and a 5% alpha error rate reflecting changes in microbial composition, based on previously observed robust differences in beta-diversity and the presence of unique taxa within each supplier-origin GM.^24,25,42^ All husbandry was performed in accordance with barrier conditions including use of autoclaved, irradiated chow, autoclaved, acidified water, and autoclaved bedding. Biweekly cage changes occurred in a laminar flow hood using bead-sterilized forceps to transfer mice between cages, by personnel wearing bleach-disinfected latex gloves. Mice were on a 14:10 light/dark cycle and were determined to be free of all known pathogens based on comprehensive quarterly sentinel testing through IDEXX BioAnalytics.

Fecal samples were collected by placing each mouse into an empty, autoclaved, microisolator cage, allowing the mouse to defecate normally, and collecting the pellet into a sterile 1 mL cryovial using an autoclaved wooden toothpick. Cryovials were then sealed and flash-frozen in liquid nitrogen. Separate fecal pellets were collected from each mouse for DNA and RNA extraction.

### DNA extraction

Fecal DNA was extracted using PowerFecal kits (Qiagen) per the manufacturer’s protocol, with the exception that the initial sample disaggregation was performed with a TissueLyser II (Qiagen), operated at 30 Hz. DNA yields were eluted in 50 µL sterile water, quantified using Qubit 2.0 fluorometer and Quant-iT dsDNA Broad Range (BR) Assay kits, and diluted to a uniform volume and concentration.

### RNA extraction

Fecal RNA was extracted using MagMAX mirVana Total RNA Isolation kits (Thermo Fisher) per the manufacturer’s protocol. RNA yields were eluted in 50 µL sterile water, quantified using Qubit 2.0 fluorometer and Quant-iT RNA Broad Range (BR) Assay kits, and diluted to a uniform volume and concentration.

### Metagenomic library preparation

Metagenomic libraries were generated from genomic DNA (250 ng) per manufacturer’s protocol with reagents supplied in the Illumina DNA Prep, Tagmentation Kit. The sample concentration was determined using the Qubit dsDNA high-sensitivity (HS) assay kit. Genomic DNA was fragmented, and short adapter sequences ligated to the ends by bead-link transposomes. Tagmented DNA was amplified using a minimum number of PCR cycles (5) to complete adapter sequences required for cluster generation and the addition of unique dual indexes. Final libraries were purified by addition of Axyprep Mag PCR Clean-up beads. The final construct of each purified library was evaluated using the Fragment Analyzer, quantified using the Qubit HS dsDNA assay kit, and diluted according to Illumina’s standard sequencing protocol.

### Metatranscriptomic library preparation

Metatranscriptomic libraries were generated from total RNA (800 ng) per manufacturer’s protocol with reagents supplied in NEBNext® rRNA Depletion Kit (Bacteria) followed by fragmentation and synthesis of cDNA using the Illumina Stranded mRNA Prep, Ligation Kit. The sample concentration was determined using the Qubit RNA high-sensitivity (HS) assay kit, and the RNA integrity checked using the Agilent Fragment Analyzer automated electrophoresis system. The rRNA was first removed from total RNA by hybridization probes using the NEBNext kit instead of poly-A RNA enrichment. The rRNA-depleted samples were then precipitated and fragmented, and double-stranded cDNA was generated from fragmented RNA, and short adapter sequences ligated to the ends. The cDNA constructs were amplified using a minimum number of PCR cycles (10) to complete adapter sequences required for cluster generation and the addition of unique dual indexes. Final libraries were purified by addition of Axyprep Mag PCR Clean-up beads. The final construct of each purified library was evaluated using the Fragment Analyzer, quantified using the Qubit HS dsDNA assay kit, and diluted according to Illumina’s standard sequencing protocol. Paired-end 150 base pair length reads were sequences using an Illumina NovaSeq 6000 instrument. All six metagenomic and six metatranscriptomic libraries were pooled to yield approximately 40 Gb per metagenomic library and 190 million paired end reads per metatranscriptome library.

### Meta-omic preprocessing, assembly, binning, and analyses

For processing metagenomic sequence data, we used the Integrated Meta-omic Pipeline (IMP) workflow^68^ to process paired forward and reverse reads using version 3.0 (commit# 9672c874; available at https://git-r3lab.uni.lu/IMP/imp3). IMP includes pre-processing, assembly, genome reconstructions and additional functional analysis of genes based on custom databases in a reproducible manner. Briefly, adapter trimming is followed by filtering the reads against the mouse reference genome (GRCm38, https://www.ncbi.nlm.nih.gov/assembly/GCF_000001635.20/) to remove any reads mapping to the host, i.e. mice. Thereafter, an iterative co-assembly of both the metagenomic and metatranscriptomic reads using MEGAHIT v1.2.9^69^ is performed. Concurrently, MetaBAT2 v2.12.1,^70^ MaxBin2 v2.2.7,^71^ and binny^72^ were used for binning the assembly, for reconstructing metagenome-assembled genomes (MAGs). Upon completion of binning, we used DASTool^73^ to select a non-redundant set of MAGs using a recommended threshold score of 0.7. Furthermore, CheckM v1.1.3^74^ was used to assess the quality of the MAGs, and the GTDB-tooklit^75^ was used to assign the taxonomy per MAG. To estimate the overall abundances of eukaryotes, EUKulele v1.0.5^76^ was used on the assemblies, with both the MMETSP and the PhyloDB databases. Each of the databases were run separately to confirm the detected eukaryotic profiles, whereby conflicts in assigned taxonomy were resolved by selecting the best hit score. To understand the overall metabolic and functional potential of the metagenome and reconstructed MAGs we used MANTIS^77^ which annotates both assemblies and MAGs alike using several databases such as KEGG,^34,35^ PFAM,^36^ and CAZyme.^37^ All the parameters, databases, and relevant code for the analyses described above are openly available at https://github.com/susheelbhanu/mice_multiomics_mmrrc and included in the Code availability section.

### Phylogenomics, pangenome construction and differential analyses

To perform the pangenome analyses, bins with the same level of taxonomic resolution, i.e., genus or family level, were collected. They were subsequently subjected to the pangenome workflow as described here http://merenlab.org/2016/11/08/pangenomics-v2, by Meren *et al. w*ithin the anvi’o^78^ ecosystem. For the *Saccharibacteria* pangenome analysis, two existing genomes (accession IDs: CP040003 and CP040004.1) from Genbank were downloaded. The pangenome was run using the -- min-bit 0.5, --mcl-inflation 10 and --min-occurence 2 parameters, excluding the partial gene calls. A phylogenomic tree was built using MUSCLE v3.8.1551^79^ and FastTree2 v2.1.10^80^ on all single-copy gene clusters in the pangenome that were present in at least 30 genomes and had a functional homogeneity index below 0.9, and geometric homogeneity index above 0.9. The phylogenomic tree was used to order the genomes, the frequency of gene clusters (GC) to order the GC dendrogram. For the Saccharibacteria phylogenetic tree, we used the *Entrez Direct* tools available at https://www.ncbi.nlm.nih.gov/books/NBK179288/, to fetch all genomes labelled as ‘Saccharibacteria’, within NCBI. Following this, the genomes were input to GToTree v1.5.51130^81^ pipeline with the -D parameter, allowing to retrieve taxonomic information for the NCBI accessions, where the tree was built using ‘Bacteria and Archaea’ marker genes. Briefly, HMMER3 v3.3.2^82^ was used to retrieve the single-copy genes after gene-calling with Prodigal v2.6.3^83^ and aligned using TrimAl v1.4.rev15.^84^ The entire workflow is based on GNU Parallel v20210222134.

### Data analyses and figures

The heatmaps were generated using the ggplot2 package while the volcano plots were built using the EnhancedVolcano package found at https://github.com/kevinblighe/EnhancedVolcano. The correlation matrices were generated using the corrplot package. Furthermore, for the differential analyses, we used DESeq2^38^ with FDR and multiple-testing adjustments to assess enriched KOs, pathways, and expression levels. For the Saccharibacteria tree visualization the following packages from the R environment were used: ape, ggree, ggtreeExtra and treeio.

## Data Availability

Raw sequencing data samples and the MAGs are available at NCBI’s sequence read archive under BioProject accession PRJNA876568. The BioSample accession IDs and the metadata associated with each sample are listed in **Supplementary Table 2**.

## Code Availability

The detailed code for the downstream analyses including the assemblies using IMP is available at https://github.com/susheelbhanu/mice_multiomics_mmrrc. The code used to generate the Figure 5 and Supplemental Figures 1-3 is available at https://github.com/ericsson-lab/metaG_metaT.

## Figure and Table Legends

**Supplemental Figure 1.**
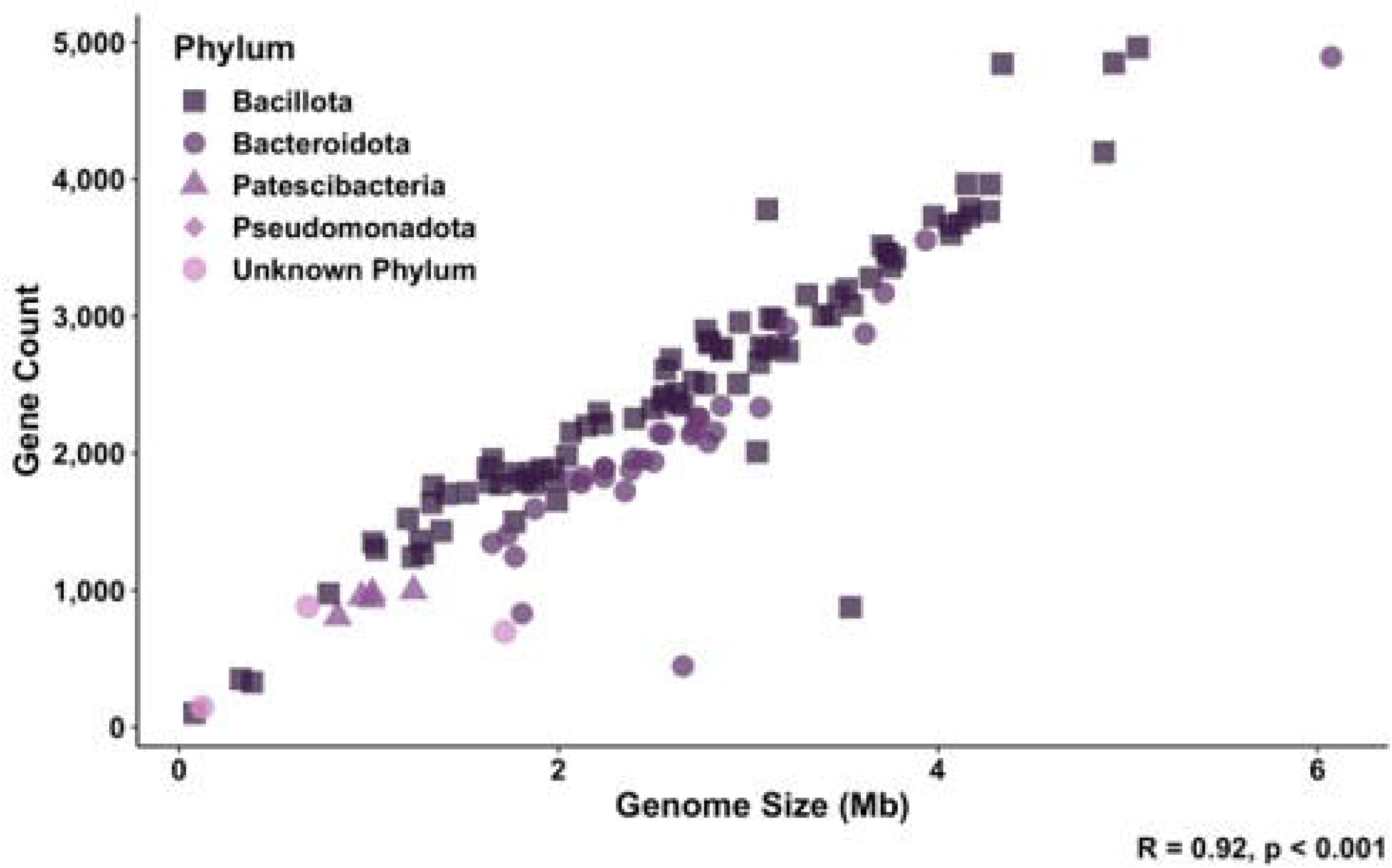
Dot plot representing the significant correlation between the number of detected genes and assembled MAG size (Mb). Spearman correlation. R = 0.92, *p* < 0.001.

**Supplementary Figure 2.**
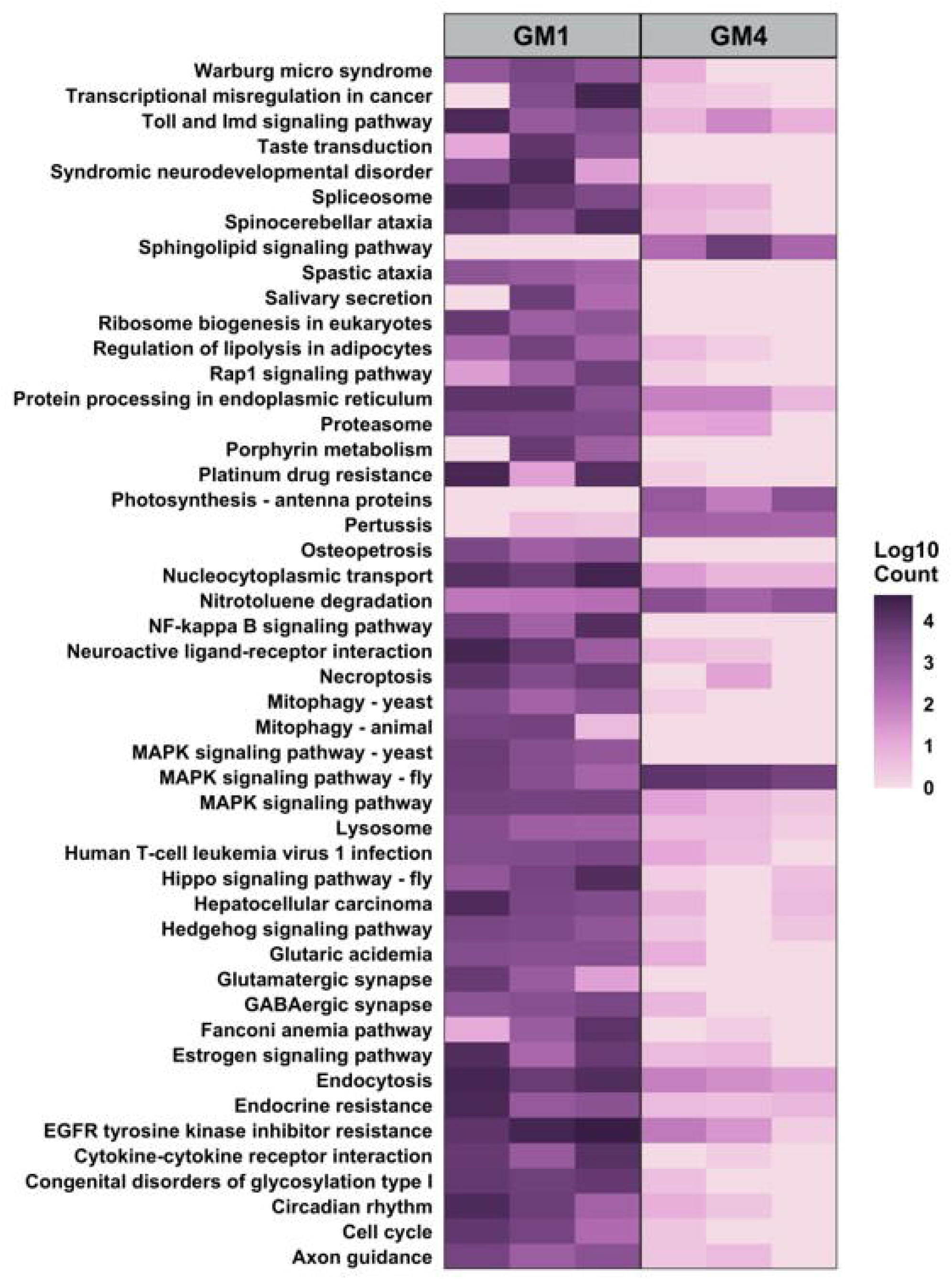
Heatmaps of host-associated pathways differentially expressed in fecal metatranscriptomic data in Jackson (GM1)- and Envigo (GM4)-origin microbiomes.

**Supplementary Figure 3.**
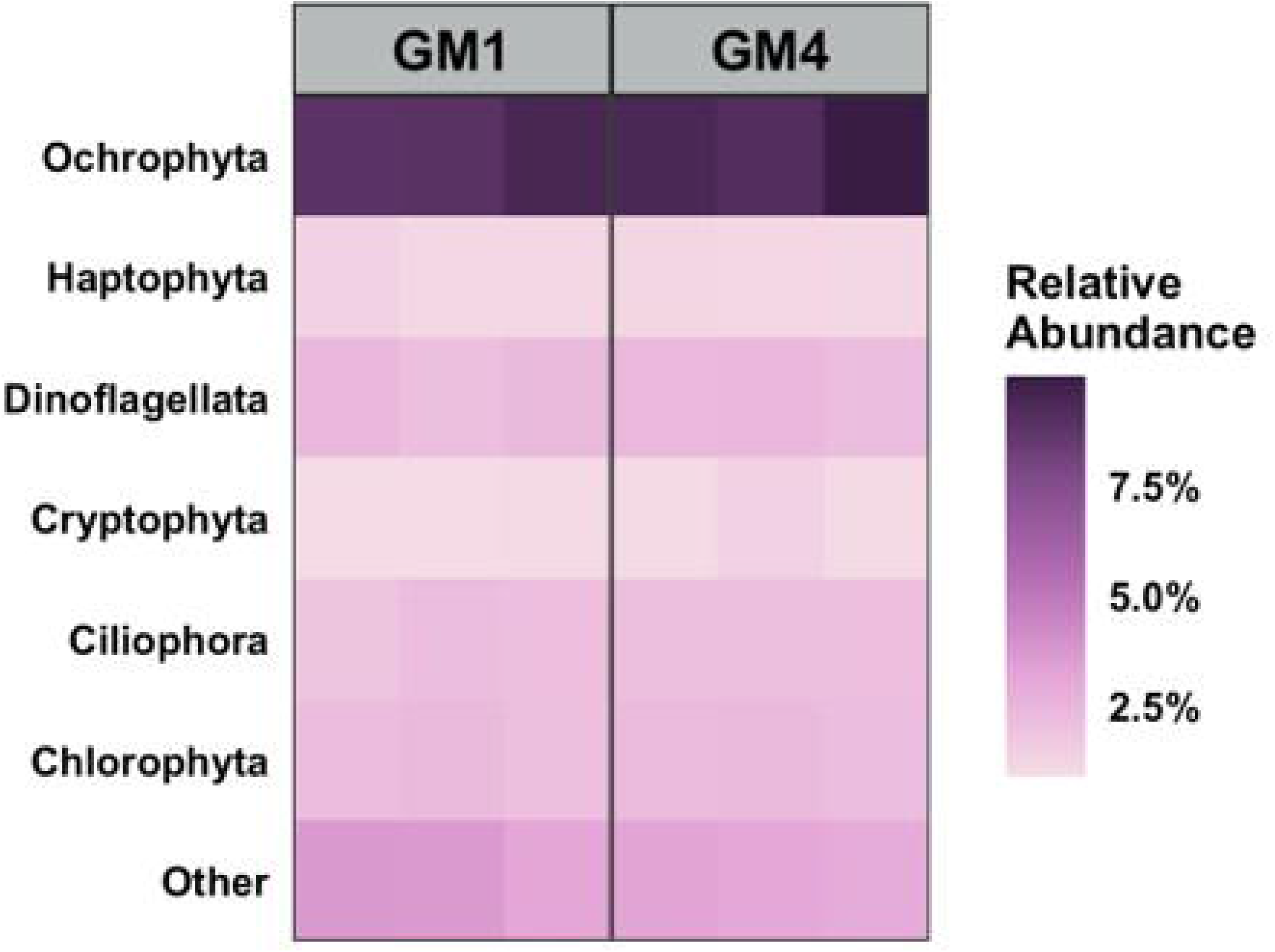
Relative abundance heatmaps of phyla representing greater than 1% of eukaryotes in Jackson (GM1)- and Envigo (GM4)-origin microbiomes.

**Supplementary Figure 4.**
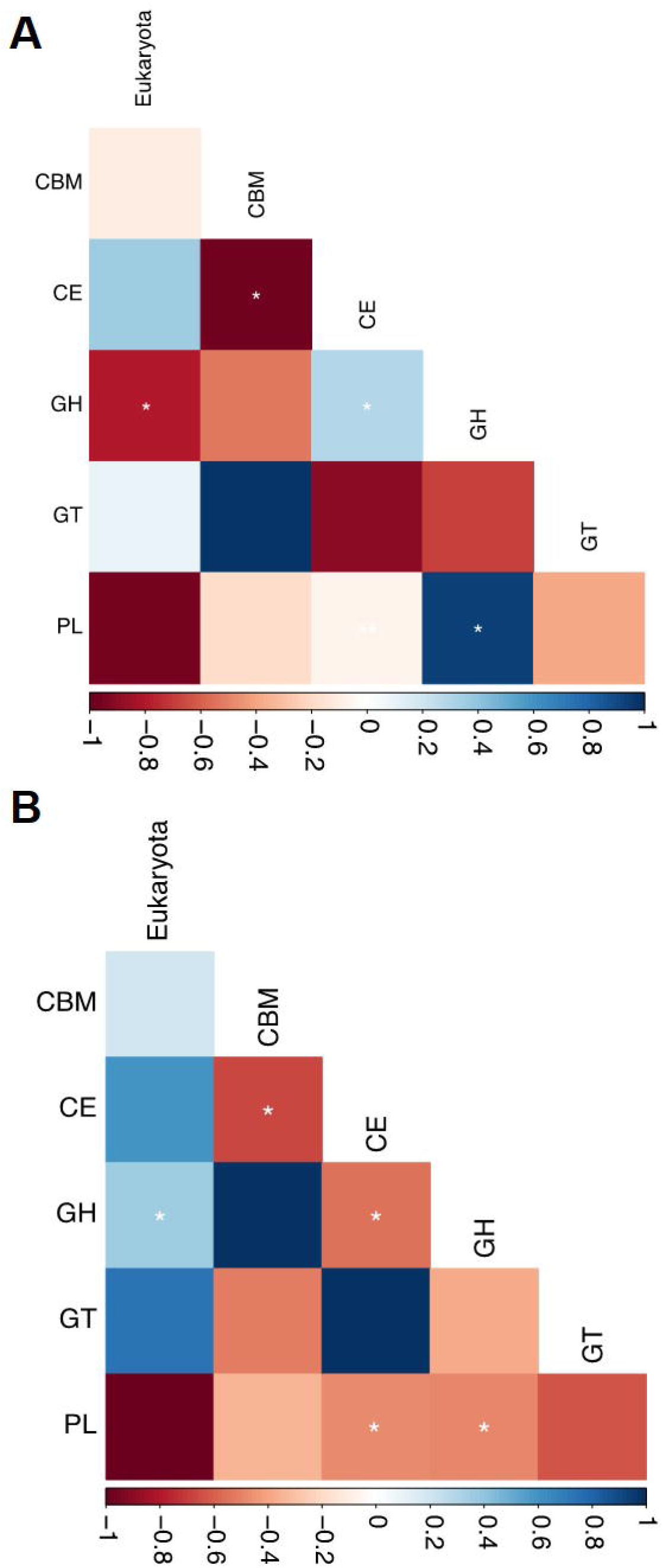
Heatmaps demonstrating correlations between classes of CAZyme molecules and detected eukaryotes in GM1 **(A)** and GM4 **(B)**. *: *p* < 0.05.

Table 1. High quality MAGs (>90% completion and < 5% contamination) identified in GM1- and GM4-origin gut microbiomes.

